# Developmental toxicity of pre-production plastic pellets affects a large swathe of invertebrate taxa

**DOI:** 10.1101/2024.01.10.574808

**Authors:** Eva Jimenez-Guri, Periklis Paganos, Claudia La Vecchia, Giovanni Annona, Filomena Caccavale, Maria Dolores Molina, Alfonso Ferrández-Roldán, Rory Daniel Donnellan, Federica Salatiello, Adam Johnstone, Maria Concetta Eliso, Antonietta Spagnuolo, Cristian Cañestro, Ricard Albalat, José María Martín-Durán, Elizabeth A. Williams, Enrico D’Aniello, Maria Ina Arnone

## Abstract

Microplastics pose risks to marine organisms through ingestion, entanglement, and as carriers of toxic additives and environmental pollutants. Plastic pre-production pellet leachates have been shown to affect the development of sea urchins and, to some extent, mussels. The extent of those developmental effects on other animal phyla remains unknown. Here, we test the toxicity of environmental mixed nurdle samples and new PVC pellets for the embryonic development or asexual reproduction by regeneration of animals from all the major animal superphyla (Lophotrochozoa, Ecdysozoa, Deuterostomia and Cnidaria). Our results show diverse, concentration-dependent impacts in all the species sampled for new pellets, and for molluscs and deuterostomes for environmental samples. Embryo axial formation, cell specification and, specially, morphogenesis seem to be the main processes affected by plastic leachate exposure. Our study serves as a proof of principle for the potentially catastrophic effects that increasing plastic concentrations in the oceans and other ecosystems can have across animal populations from all major animal superphyla.

## 1. Introduction

Plastic contamination has emerged as a significant concern in marine ecosystems due to its pervasive presence and potential impact (Eriksen et al., 2014; Thushari & Senevirathna, 2020). Large plastic items can entangle marine animals, cause physical injuries, and alter or disrupt habitats. In time, with the physical and chemical actions of the environment, plastics can break down and produce secondary microplastics, particles smaller than 5 mm. Other microplastics already arrive as small particles to the environment, known as primary microplastics. Whichever the origin, microplastics possess certain characteristics that make them a particular concern: they have a global distribution, as they are easy to disperse because of their small size; they can be ingested by a wide range of animals, entering the food chain at any trophic level; and they have a relatively larger surface area compared to larger plastic items, allowing them to adsorb, accumulate and carry pollutants from the surrounding environment (Mato et al., 2001). These pollutants can include toxic chemicals and heavy metals, leading to potential risks when ingested by marine organisms, as well as the potential to release these harmful additives into the water, creating additional hazards to marine life and ecosystems (Teuten et al., 2009; Engler, 2012; Gauquie et al., 2015; Rendell-Bhatti et al., 2021). Among the significant contributors to primary microplastic pollution, in terms of weight, are pre-production plastic pellets commonly known as nurdles (Sherrington, 2016), the building blocks for plastic products. During manufacturing, these nurdles are combined with various chemical compounds, including plasticizers, stabilizers, and antioxidants, necessary to impart specific physical properties to the final products. These chemical compounds are readily transferred to the water (Mato et al., 2001; Rendell-Bhatti et al., 2021; Paganos et al., 2023) in the case of the nurdles being lost at sea (Sewwandi et al., 2022). Once in the water, they have the ability to concentrate environmental pollutants and transport and release them to a different location, often far from the source of contamination (Teuten et al., 2009).

Many marine invertebrates commonly undergo embryonic development in the water column, and their larvae are usually planktonic. Their developmental strategy, combined with the absence of a protective eggshell, make them vulnerable to contamination from plastic leachates. Plastic leachate toxicity has been shown to have negative effects on the development of several marine organisms (Li et al., 2016; Oliviero et al., 2019; Gardon et al., 2020), and in particular, plastic pre-production pellet leachates can disrupt the development of sea urchins (Nobre et al., 2015; Rendell-Bhatti et al., 2021; Paganos et al., 2023) and to some extent brown mussels (Gandara e Silva et al., 2016). However, there is a lack of knowledge regarding how universal the susceptibility to this contamination is across other animal groups.

Here, we provide a systematic characterisation of the phenotypic abnormalities found during development and asexual reproduction by regeneration in an array of aquatic invertebrates treated with both environmental and industrial pre-production plastic pellet leachates. We select representative species of the aquatic ecosystem, including one mollusc, two annelids, one flatworm, one crustacean arthropod, two echinoderms, two tunicates and one cnidarian, to investigate microplastic-induced abnormalities in organisms across all major animal superphyla (Figure 1).

**Figure 1.**
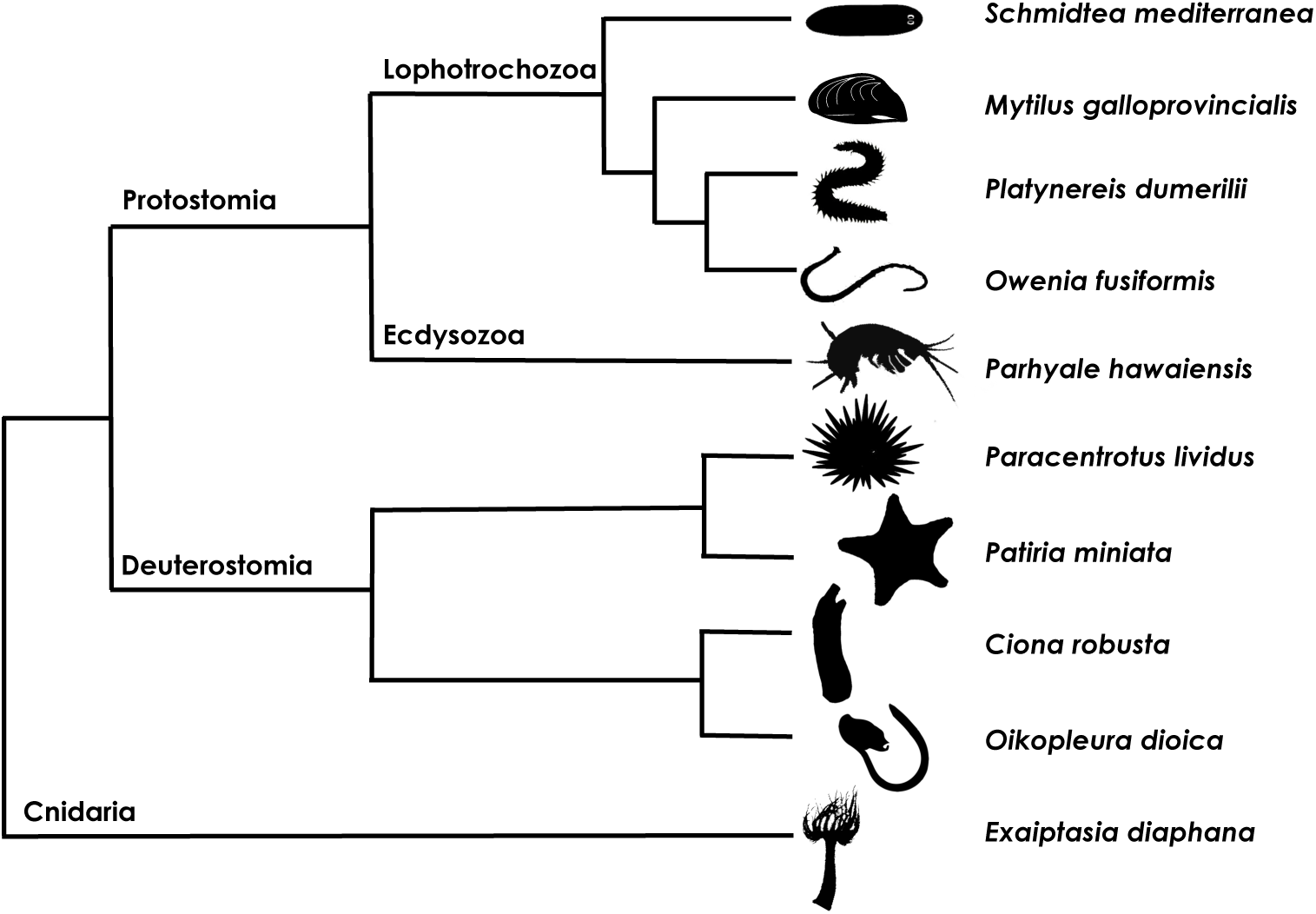
Phylogenetic tree. A schematic representation of the phylogenetic relationships among the species studied in this work. Representatives of the three major bilateria superphyla are used (Protostomia (Lophotrochozoa and Ecdysozoa) and Deuterostomia), as well as a representative of the radiata (Cnidaria). *O. dioica* by Josep Martí-Solans (CC0 1.0).

## 2. Materials and methods

### 2.1. Microplastic leachate preparation

Environmentally retrieved nurdles were obtained from Watergate bay (Cornwall, UK) by Beach Guardian CIC in December 2021 and manually sorted from other plastic particles. Commercial PVC nurdles were purchased from Northern Polymers and Plastics Ltd. (UK) in January 2022. In brief, for each of the plastic particles, pellets were added to filtered seawater (0.22 μm) (FSW) at a concentration of 10% (v/v). Pellets were leached for 72 h on a platform shaker, with continuous shaking at room temperature (ca 18°C). Leachates were obtained by filtering through filter paper in order to remove particles. Leachates were diluted to the final concentration in FSW. Only for *Schmidtea mediterranea*, leachates were obtained and diluted in Planarian Artificial Medium (1X PAM) (Cebrià & Newmark, 2005) rather than FSW. We tested 10% nurdle leachates and 1%, 5% and 10% PVC leachates (v/v), concentrations which had previously been shown to induce aberrant phenotypes in echinoderms (Rendell-Bhatti et al., 2021; Paganos et al., 2023), to produce comparable results with our previous work. Tests with lower concentrations than the ones used here did not produce evident aberrant phenotypes.

### 2.2. Animal husbandry, fertilisation and embryo exposure

Sexually mature specimens of *Mytilus galloprovincialis* were obtained from Irsvem Srl, a commercial shellfish farm (Bacoli, Napoli, Italy). Animals were mechanically stimulated to promote spawning by scraping the shells to remove adherent organisms and pulling the byssus. Approximately, 20–30 mussels were placed in a tank with Mediterranean FSW at 18°C and spread to easily monitor the spawning. When mussels began to spawn, each individual was washed and then transferred into a beaker containing 200 ml of Mediterranean FSW to isolate male and female gametes. Eggs were fertilised with an egg/sperm ratio 1:15 in a volume of 50 ml. The resulting zygotes were left to grow at 18°C in treatment plates at a concentration of 250 eggs per ml until the developmental stage of interest, the D-larva at 48 hours post fertilisazion (hpf).

*Platynereis dumerilii* gametes were obtained from an in-house culture at the University of Exeter, (UK), with culture conditions based on (Hauenschild & Fischer, 1969). Batches of embryos were created by allowing a single male and female epitoke to freely spawn in a 100ml glass beaker with 80ml 0.22 µm filtered artificial seawater. Developing eggs were stripped of their protective jelly by thorough rinsing through a 100 µm filter mesh cup one hour after fertilisation. De-jellied fertilised eggs were then added to the different treatments and left to develop in an incubator at a constant temperature of 18°C with a regular light-dark cycle (16h light, 8h dark) for 96 hours.

*Parhyale hawaiensis* were housed in artificial sea water at 24°C at the University of Exeter, Penryn campus (UK). Sexually mature pairs in amplexus were transferred to beakers containing the different treatments. When the female moults, pairs separate and oocytes are fertilised and deposited in the female’s ventral pouch (Rehm et al., 2009). After a few days, females were anesthetised with clove oil (Rehm et al., 2009) and eggs removed and transferred to containers with the same treatment as the parents. This was done to avoid predation of eggs or hatched larvae by the adults. Because of the way *P. hawaiensis* embryos are fertilised, embryos from every pairing were only assigned to one treatment, differing from the other species here studied. Embryos were left to develop at 24°C until hatching occurred.

*Paracentrotus lividus* were housed in circulating seawater aquaria at 18°C in the aquarium facility of the Stazione Zoologica Anton Dohrn, Naples (Italy). Gamete acquisition and fertilisations were performed as described elsewhere (Rendell-Bhatti et al., 2021). Embryos were added to treatment beakers immediately after confirmation of fertilisation at a concentration of 50 embryos per ml and left to develop at 18°C until the 48 hpf pluteus stage.

*Patiria miniata* were housed in circulating seawater at 14 °C, to prevent them from spontaneous spawning, in the aquarium facility of the Stazione Zoologica Anton Dohrn, Naples (Italy). Gametes were retrieved from adults by arm incision and extraction of intact gonads. Eggs were released from the ovaries through manual dissection, while male gonads were placed dry in an Eppendorf tube at 4°C. Immature oocytes were incubated with 1:1000 dilution of 10mM 1-Methyladenine in FSW for 1h at 15°C. Once the germinal vesicle was broken down, an indication of the oocytes’ maturity, they were fertilised with a few drops of diluted sperm in FSW (1μl of dry sperm in 10ml of FSW). Embryos were then added to treatment beakers at a concentration of 50 embryos per ml and left to develop at 15 °C until 4 days post fertilisation (dpf, bipinnaria larvae stage) in 9:1 diluted FSW (9-parts Mediterranean FSW, 1-part distilled water) to obtain the appropriate salinity of approximately 35ppt.

*Ciona robusta* were collected in Taranto (Italy) in March 2023, and left for at least seven days in an aquarium facility at 16-18°C with permanent light to promote gametogenesis at the Stazione Zoologica Anton Dohrn, Naples (Italy). Gametes were obtained as described before (Eliso et al., 2020), with modifications. In brief, oocytes and sperm were harvested from each individual by dissecting the gonoducts sequentially, to avoid self fertilisation. Fertilization was performed by adding diluted sperm (1:100 in FNSW) to the eggs suspension. After 15 minutes of incubation on a rotating shaker, the fertilized eggs were rinsed in FSW to avoid polyspermy and added to treatment plates about 30 minutes post-fertilization, with a density of 10 embryos per ml and left to develop at 18°C until the desired developmental stage (hatched larva).

*Oikopleura dioica* specimens were cultured in the animal facility of the University of Barcelona (Spain) as previously described (Martí-Solans et al., 2015). Mature females and males were collected separately at day 5, and left to spawn naturally. For each experiment, multiple egg and sperm batches were mixed, *in vitro* fertilised and transferred before the first cell division to treatment plates at 19°C until the desired larval stage.

*Owenia fusiformis* collected from the coasts near the Station Biologique de Roscoff were maintained in artificial seawater at the Queen Mary University of London, London (UK) at 15°C. Animals were removed from their sand tubes as described elsewhere (Carrillo-Baltodano et al., 2021) and decapitated with a razor blade just above the first parapodia before being left to regenerate in seawater supplemented with penicillin (100U/ml) and streptomycin (200 µg/ml) at 19°C.

*Schmidtea mediterranea* from an asexual clonal line were housed at 20°C in 1X PAM water (Cebrià & Newmark, 2005) at the University of Barcelona (Spain). Animals were fed twice per week with organic veal liver and were starved for at least 1 week before experiments. Pre-pharyngeal amputation was performed using a razor blade. After amputation, animals were immediately transferred to treatment plates at 20°C and left to regenerate until observations at five and seven days.

*Exaiptasia diaphana* (formerly *Aiptasia pallida*) polyp specimens were housed in circulating seawater aquarium facility at the Stazione Zoologica Anton Dohrn, Naples (Italy). For our experimental purposes, they were kept starved in crystallizing dishes at 24°C in a light/dark cycle of 12/12 hours. Pedal lacerations were collected manually and placed in a 12-well plate, one laceration per well. They were allowed to regenerate in FSW for one week. After that time, FSW was replaced with leachate solution at the defined concentration, and regenerating fragments were incubated at 18°C for one week.

### 2.3. Phenotypical observations

Larvae of the species object of the study were arrested by fixing them in 4% PFA and imaged using either a Leica M165C with a Leica DFC295 camera, or a Leica DMi8 with a Leica flexacam C3 microscope. *O. dioica*, *S. mediterranea* and *E. diaphana* were imaged alive using an Olympus SZX16 stereomicroscope, an sCM EX-3 high end digital microscope camera (DC.3000s, Visual Inspection Technology) and a Zeiss AXIO Zoom V16 microscope equipped with Axiocam 208 colour camera, respectively. Larvae were classified into two groups: normal developed larvae and aberrant larvae, including phenotypes ranging from delayed to totally aberrant. However, notes were made in the phenotypes that, despite looking normal, were not quite like the controls, as well as for delayed larvae that looked otherwise correct. Statistical differences were analysed by performing One-Way ANOVA followed by Post Hoc Tukey HSD. Mussel area size differences between controls and treated animals for each individual spawning (n=6 spawning events) were calculated with ImageJ and differences were analysed by performing unpaired t-tests.

### 2.4 *S. mediterranea* immunohistochemistry and cell proliferation counts

Whole-mount immunohistochemistry in planarians was performed as previously described (Ross et al., 2015; Fraguas et al., 2021). The following antibodies were used: mouse anti-SYNAPSIN, used as pan-neural marker (anti-SYNORF1, Developmental Studies Hybridoma Bank, Iowa City, IA, USA) diluted 1:50; mouse anti-VC1(Sakai et al., 2000), specific for planarian photosensitive cells (anti-arrestin, kindly provided by H. Orii and Professor K. Watanabe) diluted 1:15000; rabbit anti-phospho-histone H3 (Ser10) to detect cells at the G2/M phase of cell cycle (H3P, Cell Signaling Technology) diluted 1:300. The secondary antibodies Alexa 488-conjugated goat anti-mouse and Alexa-568-conjugated goat anti-rabbit (Molecular Probes, Waltham, MA, USA) were diluted 1:400 and 1:1000, respectively. Samples were mounted in 70% glycerol before imaging. Fixed and stained animals were observed with a Leica MZ16F stereomicroscope and imaged with a ProgRes C3 camera (Jenoptik, Jena, TH, Germany). Confocal images were obtained with a Zeiss LSM 880 confocal microscope (Zeiss, Oberkochen, Germany). Image processing and quantifications were performed with Adobe Photoshop and ImageJ2. Counting of the H3P-positive cells was carried out manually and normalized by the total body area. Statistical analyses and graphical representations were performed using GraphPad Prism 9. A box plot displaying the minimum, lower first quartile, median, upper third quartile, and maximum values was used to represent the data. Kruskal-Wallis test was performed to compare the means between conditions after discarding data normality and homogeneity for some samples using the Shapiro-Wilk test.

## 3. Results

### Plastic pellet leachates affect a large swathe of animal phyla

We tested the effects of leachates of high concentrations of new and beach plastic pellets on the development of *Mytilus galloprovincialis* (mollusc), *Platynereis dumerilii* (annelid), *Parhyale hawaiensis* (arthropod), *Paracentrotus lividus* and *Pariria miniata* (echinoderms), and *Ciona robusta* and *Oikopleura dioica* (tunicates) (Figure 2), and on the regeneration capacity of *Owenia fusiformis* (annelid), *Schmidtea mediterranea* (platyhelminth) and *Exaiptasia diaphana* (cnidaria) (Figure 3). Our results showed that the effects were treatment and dose-dependent, as well as species-specific (Figure 4).

**Figure 2.**
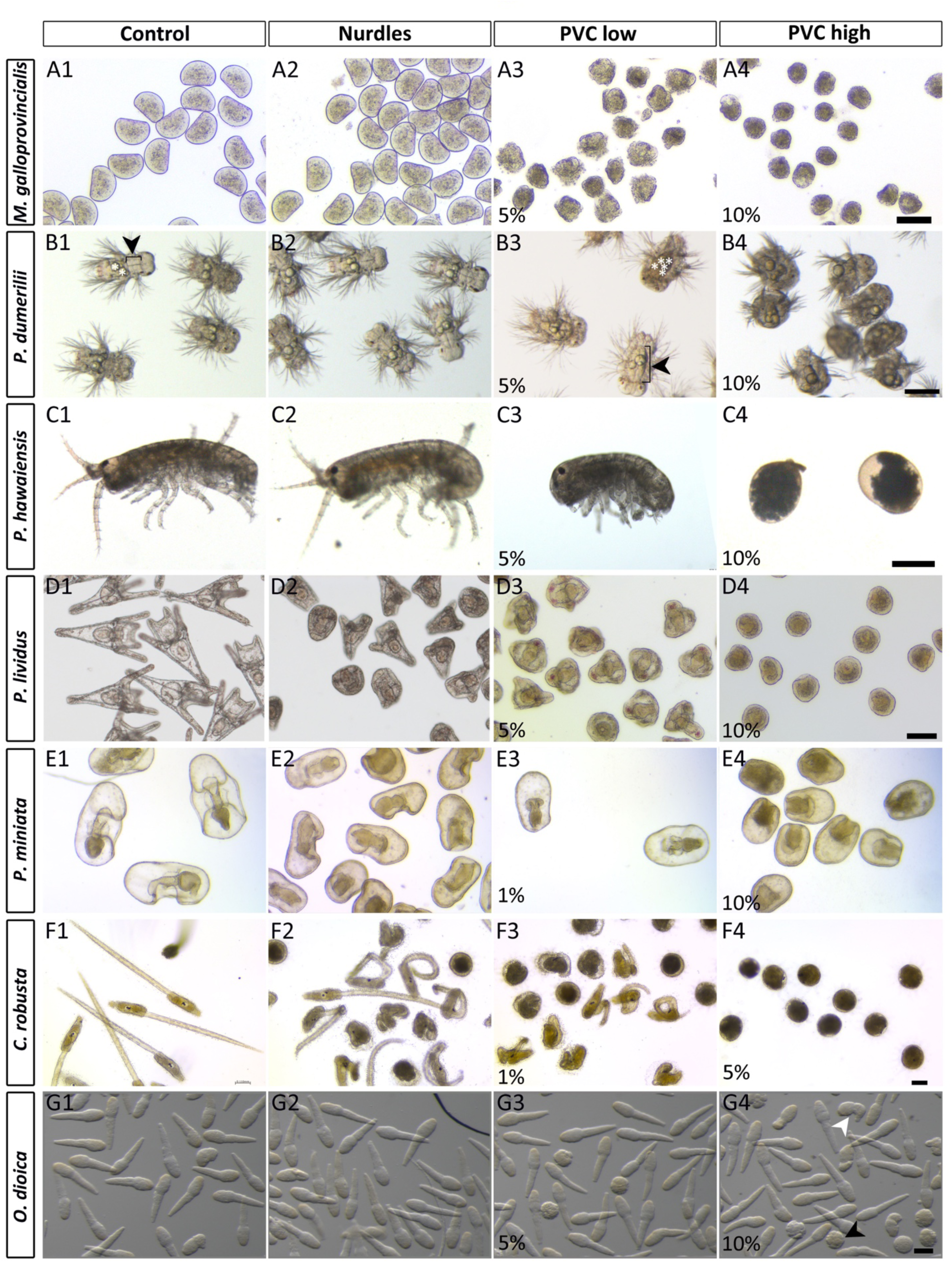
Morphological effects on larval development. Representative pictures of the phenotypes observed for every studied species as they undergo each treatment. See text for details. Columns are different treatments: (1) Control; (2) Nurdle; (3) PVC low, represents the lowest concentration tested (between 1% and 5% PVC) resulting in a significant phenotype; (4) PVC high, represents the concentration at which the most aberrant phenotype was obtained (10% PVC for all species except for *C. robusta*, which was already at 5%). Lines are different species, as shown in left panel. (B1) White asterisks show and example of normal number of lipid droplets; black bracket and arrowhead show an example of normal foregut. (B3) White asterisks show an example of abnormal number of lipid droplets; black bracket and arrowhead show an example of an enlarged foregut (G4) White arrowhead points to a kinked-tail phenotype, black arrowhead points to a golf ball phenotype. Scale bars are 110 μm for all animals except *P. hawaiensis* which represents 200 μm.

**Figure 3.**
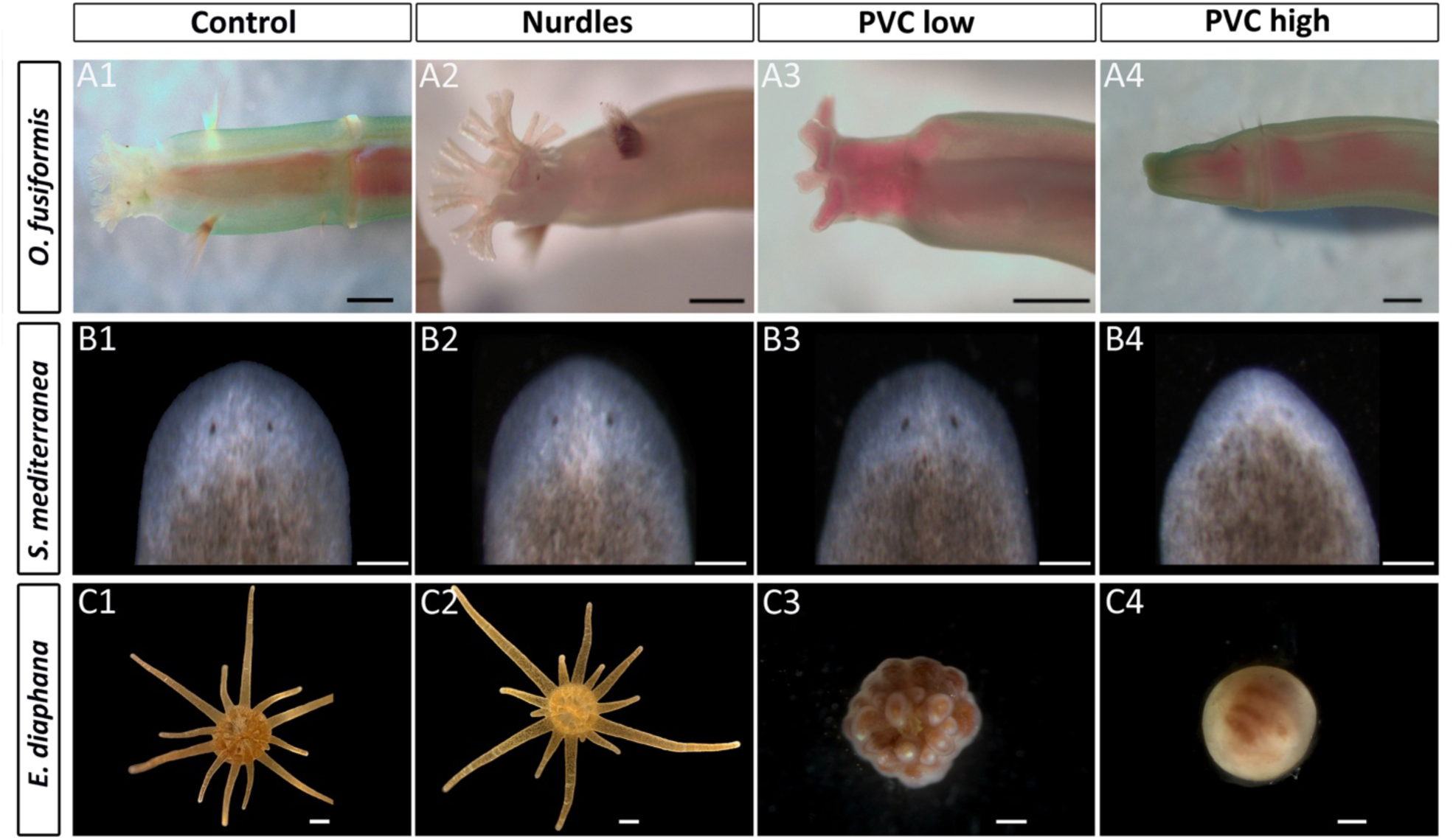
Morphological effects on regeneration. Representative pictures of the phenotypes observed for every studied species as they undergo each treatment. See text for details. Columns are different treatments: (1) Control; (2) Nurdle, 10% nurdle leachates; (3) PVC low, 5% PVC leachates; (4) PVC high, 10% PVC. Lines are different species, as shown in left panel. A and B show only the anterior regenerating heads of the animal. A, anterior to the left; B, anterior to the top. Scale bars represent 500 μm for *O. fusiformis* and 200 μm for *S. mediterranea* and *E. diaphana*.

**Figure 4.**
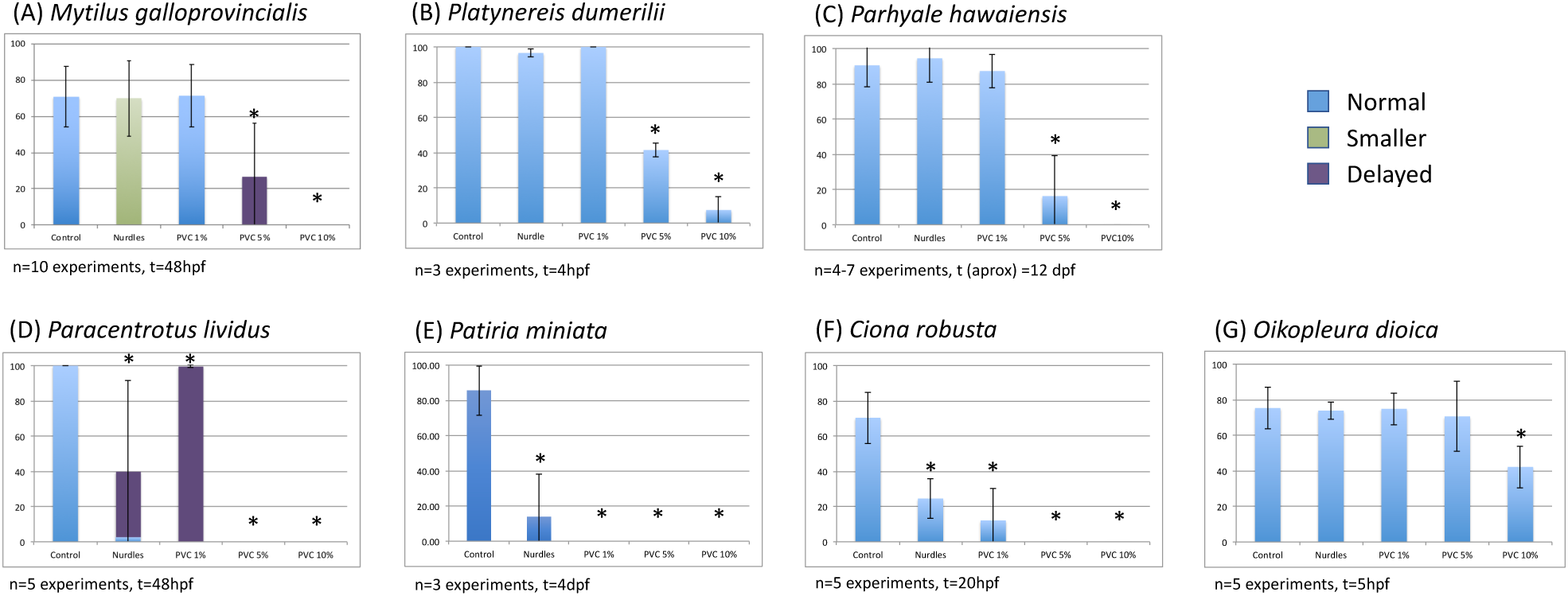
Percentage of normal developmental phenotypes obtained for each species studied in each treatment. Percentage of normal embryos is depicted in blue. When embryos are delayed but otherwise normal they are depicted in purple. Smaller than normal but otherwise normal embryos are shown in green. Number of biological replicas and time of observation are stated under each species graph. Error bars show standard deviations. Asterisks indicate differences from the controls have statistic significance (One-Way ANOVA followed by Post Hoc Tukey HSD).

We observed *M. galloprovincialis* larvae at 48 hpf. At this stage, the controls displayed normal D-larvae phenotype, with a well-formed early shell that covered the mantle of the larvae (Marin, Le Roy & Marie, 2012) (Figure 2. A1; Supplementary figure 1. A). Nurdle-leachate-treated larvae had a very similar phenotype to the controls, but they were slightly smaller in size (p<0.05) (Figure 2. A2; Supplementary figure 1. A, B, F). No differences were detected between 1% PVC-leachate treated embryos and controls (Supplementary Figure 1. C), but at 5% there was a significative reduction in size, with a misshaped shell in what we classified as an aberrant larva with protruding mantle (Figure 2. A3; Supplementary figure 1. D). At 10% PVC-leachate, the larvae did not develop properly and remained arrested around the trochophore stage, barely forming, in some instances, a very rudimentary incipient shell (Figure 2. A4; Supplementary figure 1. E).

*Platynereis dumerilii* develops first into a trochophore larvae, followed by a nectochaete larval stage (Özpolat et al., 2021). We assessed morphological changes at 4 dpf, the nectochaete larva. At this point, control and nurdle-treated larvae looked the same, with normal segmentation, chaetae, digestive system and lipid droplet distribution (Figure 2. B1, B2). However, larvae treated with PVC leachates did not develop properly. For 1% PVC leachate-treated larvae, despite looking otherwise normal, a few of the animals showed a deformed gut phenotype (not shown). This gut phenotype was more pronounced and common in 5% PVC-treated animals (Figure 2. B3). While the rest of the larvae could be considered normal, the developing digestive system showed a probable over-extension of the foregut tissue, and the lipid droplet distribution was also aberrant. In the 10% PVC leachate treatment, many larvae failed to complete the trochophore-to-nectochaete transition (Figure 2. B4). Where these larvae had chaetae, they showed that segmentation was not properly completed, and they resembled a truncated malformed nectochaete. The lipid droplet distribution of these larvae was also abnormal. A normal nectochaete should have four prominent lipid droplets, with two larger and two smaller droplets, sitting just below the foregut-mouth region, but these larvae showed more droplets, and oddly distributed. A normal trochophore displays four big lipid droplets (Fischer, Henrich & Arendt, 2010), and the higher number of droplets seen in the truncated nectochaete could be a remnant of the failed transition between the two larval stages. The larvae that became more like nectochaete also showed apparent problems with the developing gut similar to what is seen at 5% PVC, potentially due to an inhability to differentiate the foregut from the rest of the gut, or over-proliferation of the foregut.

*Parhyale hawaiensis* hatches into a juvenile larva after about 10 dpf. Control larvae showed a normal phenotype, with a normal head with two antennae segments, a thorax with two claws and five legs and an abdomen (Figure 2. C1). No changes were seen in the nurdle or 1% PVC treatments (Figure 2. C2; not shown). For 5% PVC leachate-treated animals, about half of the larvae showed malformed appendages, both at the level of the head, where the antennae formed but were deviated toward the posterior of the head, and the thorax, where claws and legs were malformed and twisted (Figure 2. C3). None of the 10% PVC leachate-treated embryos developed properly and all died *in ovo* (Figure 2. C4).

*Paracentrotus lividus* were imaged at 48 hpf, at pluteus larva stage. Control larvae are bilaterally symmetrical four-arm pluteus, with the typical tripartite gut, ciliary band and skeletal rods (Figure 2. D1). Nurdle leachate-treated larvae are either delayed or malformed (Figure 2. D2; Figure 4. D). These malformations include shorter or absent arms, probably due to skeletogenic impairment and a bell-shaped morphology consistent with a radialisation problem. This phenotype agrees with the one observed before in this species (Rendell-Bhatti et al., 2021), if slightly milder, probably due to different plastic particles used (see discussion below). All PVC leachate treatment concentrations show developmental abnormalities in *P. lividus*. 1% PVC leachate-treated larvae are delayed, with no other clear phenotypic abnormality (not shown; Figure 4. D). 5% PVC leachate-treated embryos display a bell-shape, being clearly radialised. Malformation of the skeleton and of the distribution of the pigment cells is also evident (Figure 2. D3). 10% PVC-treated larvae have a very extreme phenotype. In some cases, they exhibit a radialised phenotype with no elongation of the arms and a lack of pigment cells. In other cases, the archenteron has not even elongated and, in most cases, the embryo has not proceeded post-gastrulation (Figure 2. D4).

*Patiria miniata* produce a bipinnaria larvae (Yankura et al., 2010). Control larvae at 4 dpf show a typical young bipinnarial larva phenotype, including a partitioned digestive tract, elongated coeloms that will give rise to the hydrovascular organ (Perillo et al., 2023) and a well-formed ciliary band (Figure 2. E1). All treated larvae show some delay or aberrant phenotypes. For nurdle-leachate-treated embryos, larvae resemble late gastrulae (Figure 2. E2), delayed more than 24 hours from the normal developmental milestone. In many cases the gut appears like a non-partitioned tube, accompanied by ectodermal deficits. For instance, ectodermal regions such as the ciliary band and oral hood are missing. Last but not least, the elongation of the coelom appears to be delayed. For PVC-leachate treated embryos, a concentration-dependent delay in development is seen, with animals looking like late gastrula when treated with 1% PVC-leachates (Figure 2. E3), to mid gastrula, with some misshaping of the elongating archenteron at 10% (Figure 2. E4).

*Ciona robusta* tadpole larvae have two main structures: the trunk, which contains the adhesive organ (palps), the brain vesicle with two pigmented sensory organs (otolith and ocellus), endoderm and mesenchyme; and a straight tail for locomotion, bearing the neural tube, notochord, endodermal strand and muscles, all covered by the larval tunic. Control larvae showed a normal trunk and straight tail containing vacuolated notochord cells (Figure 2. F1), while all the treatments compromised the normal embryo development. Nurdle-leachate-treated larva showed a shorter, kinked or coiled tail, often disorganized in its internal structure. In most cases, the trunk shape was abnormal, the sensory vesicle was deformed, although carrying both pigmented organs, and the adhesive organ was misshaped. (Figure 2. F2). The phenotype for 1% PVC leachate-treated larvae was very similar to the observed for nurdle leachates (Figure 2. F3), although a higher percentage of embryos did not hatch and were still in the chorion. However, at 5% PVC leachate treatments, the larvae were not formed, but instead, unhatched round individuals were obtained. There were, however, two pigmented spots, probably a hint of structures corresponding to the otolith and the ocellus, demonstrating development had proceeded, but morphogenesis had not been successful, thus producing aberrant embryos (Figure 2. F4).

*Oikopleura dioica* develops extremely fast, with hatchling larvae appearing at 3.6 hpf at 19°C and larval development lasting a further 7 hours only, when the juvenile form is ready to make the first house (Ferrández-Roldán et al., 2019). The larvae, as in *C. robusta*, consist of a trunk that will house all the organs in the adult, and the tail with the notochord, the muscle cells and the nervous system. We only observed a significative shift from the controls at the highest concentration of PVC leachates (10%) when the percentage of malformed larvae increased (Figure 2. G4, Figure 4. G). These malformations affected the tail, which appeared shorter or kinked, and the trunk, which was misshaped. We also saw a higher proportion of animals that arrested their development at a pre-tailbud stage, previously described as a golf ball phenotype (Torres-Águila et al., 2018).

*Owenia fusiformis* is capable of anterior regeneration after traumatic injury. Three days after amputation, the blastema is formed, and differentiation and regeneration are complete seven days after injury (Marilley & Thouveny, 1978). At this time point, we found no differences between controls and nurdle leachate-treated animals (Figure 3. A1, A2). Heads regenerated to create fully formed tentacles and eyes. Animals treated with 5% PVC leachates looked normal (Figure 3. A3), but showed a less elaborate branching in the crown of tentacles, and two out of six did not develop the eyes properly. Likewise, five out of six of the 10% PVC leachate-treated animals regenerated properly but the branching pattern of the tentacles looked delayed. One animal failed to undergo morphogenesis at this concentration after creating an elongated blastema (Figure 3. A4).

The asexual strain of *Schmidtea mediterranea* uses stem-cell based regeneration as its main reproductive strategy. and can regenerate new heads, tails, sides, or entire organisms from small body fragments in a process taking days to weeks (reviewed in (Reddien, 2018)). We found no differences in regeneration between controls and nurdle or 5% PVC leachate-treated animals (Figure 3. B1-B3; Supplementary Figure 2). However, animals treated with 10% PVC leachates developed a smaller blastema than the controls and, at 7 days after amputation, they have regenerated smaller eyes and brains (Figure 3. B4; Supplementary Figure 2)). To investigate the proliferative rate of stem cells, we identified the G2/M stage of the cell cycle by performing immunostaining against a phosphorylated form of Histone-3. We quantified the total number of mitoses in amputated planarians regenerating anterior wounds at 5 days after amputation and observed a significant decrease in the number of mitotic stem cells in 10% PVC leachate-treated animals, but not in any other treatment (Supplementary Figure 2).

*Exaiptasia diaphana* exhibits asexual reproduction capability, growing from pedal lacerations, a small portion of tissue deriving from the margin of the pedal disk and the body column. This produces of crescent-shaped fragments that successfully regenerate into fully formed polyps within a few weeks (Clayton & Lasker, 1985; Presnell, Wirsching & Weis, 2022) (Figure 3. C1). Fourteen days post-laceration, the morphology is adult-like, differing only for the smaller size. Treatment with nurdle leachate at a concentration of 10% did not seem to affect the external morphology, which remained comparable to the control (Figure 3. C2). At a concentration of 1% PVC, the morphology of the polyps was also equivalent to the control group (not shown). However, higher concentrations of PVC leachate displayed concentration-dependent effects on normal development. Specifically, treatment with 5% PVC leachate caused developmental delays and reduced tentacle length without affecting the overall number of tentacles (Figure 3. C3). This result was consistent with the development period (Presnell, Wirsching & Weis, 2022). Furthermore, at a concentration of 10% PVC leachate, the regeneration of *E. diaphana* was severely compromised. Despite remaining alive, the laceration was covered with a protective mucous membrane (typical of new pedal lacerations), and the animal maintained its initial state without any signs of tentacle or gut formation (Figure 3. C4).

## 4. Discussion

### Developmental susceptibility across animal superphyla

The species employed in this study showed different developmental susceptibility to the treatments (Figure 4). In summary, all species were affected by new PVC particle leachates, but only some deuterostomes and the mussels had a clear response to environmental nurdle leachates. *P. dumerilii*, *O. dioica* or *P. hawaiensis* are unaffected by nurdle leachate treatment. For PVC treatments, some species are affected already at low concentrations, like *C. robusta* and *P. miniata*, which show severe phenotypes already at 1%, while the rest show clear aberrations at 5%, except for *O. dioica* which seems to be more resistant and is only affected by the 10% treatment.

For deuterostomes, in the case of the echinoderms *P. lividus* and *P. miniata* the effect seems to start early, with problems with gastrulation and archenteron elongation, as well as embryonic axis formation, as seen in other studies for *P. lividus* and *S. purpuratus* (Rendell-Bhatti et al., 2021; Paganos et al., 2023). In agreement with the phenotypes we observe, molecular data for *S. purpuratus* treated with 10% PVC leachates had previously shown downregulation of genes involved in the secondary axis formation as well as in the cell specification of certain cell types (Paganos et al., 2023). Looking into the tunicates, *C. robusta* is affected by all treatments while *O. dioica* is more resilient, only showing aberrant phenotypes at the higher concentration of PVC. In both cases, we believe axial patterning and cell specification are correct, but that morphogenesis is altered. This is clear for the treatments that display weaker phenotypes, where structures like the trunk and the tail have formed, but the morphology of these structures is aberrant (*C. robusta*, Figure 2. F2, F3; *O. dioica*, Figure 2. G4). In the most extreme phenotypes, both species display a golf ball phenotype. In *C. robusta* it is still possible to identify the two pigmented spots within the otherwise amorphous ball phenotype. For *O. dioica,* both aberrant phenotypes observed here are also seen when this animal is exposed to diatom bloom-derived biotoxins(Torres-Águila et al., 2018). Molecular analysis of the golf ball phenotype in *O. dioica* has previously showed that cell and tissue specification in this phenotype are correct, but that it is morphogenesis that is at fault (Torres-Águila et al., 2018), and we believe this is also the case here. All protostomes tested, *M. galloprovincialis*, *P. dumerilii* and *P. hawaiensis,* display a later effect at lower PVC concentrations, since they show correct axis specification and early morphogenesis. The appendage malformation in *P. hawaiensis* and the aberrant gut formation in *P. dumerilii* nectochaete larvae and the failure to produce a shell in *M. galloprovincialis* point to later gene regulatory pathways being affected in these cases. However, high PVC concentrations affect larval metamorphosis in *P. dumerilii*, as seen with the inability to properly transit from trochophore to nectochaete larva, and *P. hawaiensis* embryos fail to complete their developmental program.

### Effects on regeneration

Regeneration is a type of asexual reproduction strategy widely adopted among aquatic invertebrates by which an animal can regrow certain body parts from just a part of the original organism. Studying regeneration is important to understand healing and repair mechanisms, including those happening in humans (Mehta & Singh, 2019). Regeneration involves three main events: wound healing, cell population mobilisation (of stem cells, dedifferentiated or transdifferentiated cells) and tissue morphogenesis. *E. diaphana* can regenerate a whole polyp from a small part of the peduncle (pedal laceration) (Clayton, 1985). In the planarian *S. mediterranea*, residing pluripotent cells called neoblasts are recruited in the wound site to generate a blastema, which will then differentiate, and morphogenetic processes will assure that the correct pattern is formed to create the missing body parts (Reddien, 2018). Following anterior amputation in *O. fusiformis*, several tissue rearrangements are in place to generate a blastema where epidermal and muscle cells proliferate (Fontés et al., 1983). In our study, we see a clear hindering of the regenerative process in *E. diaphana* treated with PVC leachates (Figure 3. A3, A4). In the case of this species, wound healing takes place before we expose the animals to the treatment. Later, we cannot discern if the regeneration is obstructed at the level of cell mobilisation or tissue morphogenesis. The regeneration defects of PVC leachates in *S. mediterranea* and *O. fusiformis* are much milder (Figure 3. B1-B4, C1-C4). While *O. fusiformis* mostly shows only a delay of the regeneration process at the highest concentration of PVC leachates, the planarian displays aberrant regeneration with the formation of a smaller blastema and regenerating smaller eyes. In this case, we were able to pinpoint a reduction in the proliferation of the stem cells as well as in the differentiation of the nervous system and the new eyes (Supplementary Figure 2). This points to a correct wound healing and cell mobilisation in the planarians, and alteration in cell proliferation being the main cause of the failure in proper regeneration in this species. Whichever the mechanisms impeded by plastic leachate treatment in each of these species, there is a hinderance in regeneration because of these treatments, showing that asexual reproduction can also be affected by microplastic contamination. Indeed, nanoplastics have been found to delay regeneration in *Girardia tigrina* (Cesarini et al., 2023a) and *Hydra vulgaris* (Cesarini et al., 2023b), but to the best of our knowledge, no report has shown effects of plastic leachates on regeneration prior to our work. However, with the species sampled here, our results suggest that regeneration is more resilient to plastic contamination than development is. Further analysis of the cellular types and gene expression after infliction of the wound will be necessary to determine if pluripotent-cell recruitment and proliferation are happening correctly and if tissue morphogenesis is affected. Knowing the effects that plastic contamination can have in the regenerative processes will be informative both for the effects in asexual reproduction in marine invertebrates and for the study of possible consequences for tissue healing and regeneration in other species, potentially including vertebrates and humans subjected to plastic exposure.

### Chemical pollutants in the water

We have previously determined the content of persistent organic pollutants (POPs) and other contaminants in the leachates of these pellets (Rendell-Bhatti et al., 2021; Paganos et al., 2023). In no case were any phthalates found in the leachates of either plastic particles (Rendell-Bhatti et al., 2021). In 10% nurdle leachates, we had previously found high concentrations of polychlorinated biphenyls (PCBs) and polycyclic aromatic hydrocarbons (PAHs), 21 and 5 times higher than normal sea water content (Rendell-Bhatti et al., 2021), which could explain the developmental effects found in *P. lividus* treated with nurdle leachates (Rendell-Bhatti et al., 2021). These chemicals had also been found in similar concentrations in leachates from farming fishing gear in French Polynesia, which also induced developmental defects in the pearl oyster (Gardon et al., 2020). However, the content of POPs in 10% PVC leachates were significantly lower, and the presence of these compounds alone could not explain the stronger developmental abnormalities seen in PVC leachate treated *P. lividus* (Rendell-Bhatti et al., 2021). Analysis of the elemental content of the water leachates by inductively coupled plasma – optical emission spectrometry revealed that 10% PVC leachates contained high amounts of zinc (1 µg g ^-1^), but nurdle leachates did not have higher concentrations of zinc than sea water (Paganos et al., 2023). The zinc present in the PVC leachates could explain the phenotypes observed with this treatment, which were consistent with classical developmental experiments exposing sea urchin embryos to heavy metals (Mitsunaga & Yasumasu, 1984; Hardin et al., 1992; Kobayashi & Okamura, 2004; Cunningham et al., 2020; Paganos et al., 2023). Since using the same plastic particles, these same chemicals are expected to be the ones responsible for the phenotypes observed in the present study.

### Intraspecific variability of the response

We observed that the impact of each leachate treatment can differ in severity across batches of animals of the same species. Indeed, experiments conducted with animals collected on different days showed more variation compared to those conducted with animals collected on the same day. These differences were particularly noticeable for the nurdle and low PVC concentration leachate treatments, which generally resulted in a lower percentage of aberrant embryos than in the higher PVC leachate treatments (see, for instance, standard deviations for each species in Figure 4). Oxidative stress is highly involved in the effects of plastic and plastic-leachate treatments in marine larvae ((Paganos et al., 2023); also reviewed in (Hu & Palić, 2020)) and adults (Jeong et al., 2017; Pérez-Albaladejo, Solé & Porte, 2020; Milito et al., 2020; Murano et al., 2023). Recently, our lab and others have shown that adult sea urchins subjected to higher oxidative stress produce less successful embryos than animals in normal physiological conditions (Masullo et al., 2021; Jimenez-Guri et al., 2023). We believe that the intraspecific differences we see here are due, probably amongst other reasons, to the state of the parents before obtaining the gametes for the experiments as well as to the influence of the different genetic backgrounds of the various parents used.

We also detected intraspecific differences due to a batch effect in the particles used. For beach-collected nurdles, we found the phenotypes observed are not always equivalent when different batches of environmental nurdles are used for one species. This is inherent to the sample type, as every environmental nurdle can have a different history from when they were lost at sea and, therefore, can have gathered different types of contaminants in their travel (Teuten et al., 2009). Likewise, different types of new plastic pellets will be supplemented with, and therefore be able to leach, different chemicals at production. Therefore, it is necessary to stress again that different types of nurdles will produce different phenotypes, and that the ones shown here are only examples of what can happen. Moreover, the concentrations of pre-production plastic pellets used in this study to create the leachates are higher than those found in the sea, except maybe in the event of a nurdle spill (like the one that occurred in Sri Lanka in 2020 (Sewwandi 22) amongst others). Still, this study constitutes a proof of principle to describe that aquatic animals from different and diverse phyla are susceptible to plastic leachates during development, metamorphosis and regeneration.

### Interspecific variability of the response

Plastic contamination has been previously shown to affect development in a variety of animals, and plastic leachates are sufficient to cause these effects (Nobre et al., 2015; Li et al., 2016; Gandara e Silva et al., 2016; Oliviero et al., 2019; Gardon et al., 2020; Rendell-Bhatti et al., 2021; Paganos et al., 2023). Micro– and nano-plastics activate oxidative damage and inflammatory responses, leading to adverse outcome pathways in a variety of organisms (Hu & Palić, 2020), while plastic additives have also been shown to induce oxidative stress in aquatic organisms (reviewed in (Pérez-Albaladejo, Solé & Porte, 2020)). Previous studies have shown that adult and developing sea urchins exposed to plastic leachates have increased oxidative stress and lower production or maintenance of immune cells (Jimenez-Guri et al., 2023; Paganos et al., 2023). Furthermore, the gene regulatory networks acting early in the development and necessary for proper embryo formation are affected in these animals too (Paganos et al., 2023). It would be of great interest to look into the mechanisms of action of these toxicants to learn how leachates affect the animals studied here. One would expect that detoxification genes can be important in protecting some of the more resilient species. In *O. dioica*, response genes against environmental stress (also known as the defensome (Goldstone et al., 2006)), particularly those dealing with aldehyde detoxification, are upregulated very early in development as a response to biotoxins, where phenocopies of the larval shapes seen in our study are obtained (Torres-Águila et al., 2018). However, *C. robusta* is a species that typically lives in contaminated environments, since it thrives in ports where oil and plastic pollution, amongst others, is high, and should therefore have a very developed detoxification system. Indeed, exposure to metals in *Ciona* is known to upregulate glutathione biosynthesis (Franchi et al., 2012) and upregulate transcription of Cu,Zn superoxide dismutases (Ferro et al., 2013) and metallothionein genes (Franchi et al., 2011), all of which processes are linked to exposure and protection against oxidative stress. However, this particular species is one of the most affected by our treatments.

Oxidative stress coping mechanisms, including increasing reduced glutathione, have also been shown to be in place in *M. galloprovincialis* reared in aquaculture environments, which have increased intake of microplastics (Capo et al., 2021), and increased metallothionein expression has also been shown in *M. galloprovincialis* and *S. purpuratus* exposed to microplastics (Paganos et al., 2023; Impellitteri et al., 2023). It would, therefore, be very interesting to study the transcriptomic state of the defensome of the species studied here to correlate the different phenotypes with alterations in these specific genes.

We hypothesised that another possible reason for the interspecific differences in the severity of the phenotypes, besides specific physiological, developmental and metabolic responses to the treatments, would be the speed of development of the different species, as well as the presence of a longer-lasting, stronger chorion, as a physical barrier to the exterior. If an animal is faster at developing, it will make sense that chemicals in the water may have less time to affect their development, as they still need to pass through the chorion before it can generate stress or affect the transcription of genes. On the contrary, it could be that a slow-developer could have more time to react activating the stress-defence response. Chorions are indeed protective barriers against polymer microsphere toxicity in zebrafish embryos (Feng et al., 2013). Here, we have a mix of species that develop extremely fast (*O. dioica* hatches just 4 hpf) to relatively slow (*P. hawaiensis* hatches around 12 dpf). We do not see any emerging pattern that would allow us to determine if developing fast or slow is advantageous against plastic leachate contamination. However, the protective function of the chorion seems not to influence the effect of our treatments: despite *C. robusta* having a very strong chorion (not only composed of a surrounding membrane but also test cells, a group of maternal cells placed between the egg and the membrane, and maternal follicle cells outside the membrane (Kourakis et al., 2021)), it showed a clear and visible impairment in the proper development of the embryos. Notwithstanding the mechanism of each species to cope or react to the plastic leachates, we believe that transcriptomic profiling and determination of the oxidative stress state of these species after treatment could shed light on the mechanisms behind the differences and similarities between them.

### Ecological consequences

The potential biological impacts of plastic pollution have been extensively discussed, including the chemical toxicity from plastic leachates (Oliviero et al., 2019; Ke et al., 2019; Shi et al., 2019; Rendell-Bhatti et al., 2021; Jimenez-Guri et al., 2023; Paganos et al., 2023). Chemical stressors from plastics have been identified as a possible contributor to biodiversity loss (MacLeod et al., 2021). Embryo development is an extremely robust process that has evolved cellular processes and regulatory pathways to respond to environmental stressors. However, some anthropogenic stressors can elude the mechanisms that act as developmental defences to ensure the right developmental decisions, overwhelming the developmental robustness (Hamdoun & Epel, 2007). Problems in the correct development of embryos can lead to declines in the success of subsequent generations for a particular species. Developmental effects with no detectable phenotypes, which may be happening in the lower concentrations tested here, could also mediate potential transgenerational changes, which in turn may be detrimental. Given that many species have a spawning season, sporadic environmental stressors such as surges in local plastic contamination could impact a particular population’s reproductive success. Here, we demonstrate that plastic leachate contamination can affect the development and regeneration of many aquatic species with potentially catastrophic effects that can result in the loss of communities and disruption of ecosystems. Our study provides data showing that microplastic-derived chemical pollution can affect major lineages of marine diversity, pointing to consequences for the health and functioning of marine ecosystems.

## 5. Conclusions

Leachates of industrial PVC pellets affect all animals tested in a concentration-dependent and species-specific manner. Leachates of environmentally retrieved nurdles at the same high concentrations have a phenotypical effect in fewer species, but still in four out of the ten species tested. Deuterostomes, excluding *O. dioica*, show the highest sensitivity to all leachates. This work constitutes proof of principle that both new and environmentally retrieved pre-production plastic pellets, at high concentrations, can release enough chemicals to affect the development and regeneration of a wide group of animals. Whether this can be generalised to other types of plastic leachates is plausible but needs to be studied. Moreover, there may be effects that do not show an evident phenotype but may be affecting development, physiology or robustness at a lower level, and cumulative effects may eventually be hindering optimal development, probably including transgenerational effects. Follow-up mechanistic studies to understand the reason for the failure of the developmental and regeneration process and any effects with no detectable phenotypes at lower concentrations would be extremely interesting to address the potential ecological consequences of plastic pollution on the marine ecosystem.

## Acknowledgements

Emily Stevens and the team of Beach Guardian for providing beach plastic pellets. Aziz Aboobaker for supplying the culture of *P. hawaiensis*, Alessia Di Donfrancesco for help with the set up of the *P. hawaiensis* culture in Exeter, and Alex Hayward for transporting the animals from Oxford to Penryn. Toby Doyle for help with *P. hawaiensis* maintenance and help with running the experiments. Davide Caramiello and Alberto Macina for animal husbandry at the Stazione Zoologica Anton Dohrn. Sophie Den Hartog for help with *P. dumerilii* husbandry in Exeter University and Anna Bawden for help with sample transport. Sebastian Artime for assistance with the *O. dioica* facility in Barcelona. Francesc Cebrià for reagents towards the *S. mediterranea* experiments, and Hidefumi Orii and Kenju Watanabe for providing anti-VC-1 antibody. Monoclonal anti-SYNORF1 antibody was obtained from the Developmental Studies Hybridoma Bank, developed under the auspices of the National Institute of Child Health and Human Development and maintained by the Department of Biological Sciences, University of Iowa, Iowa City, IA, USA.

