## Supplementary figures for "Developmental toxicity of pre-production plastic pellets affects a large swathe of invertebrate taxa"

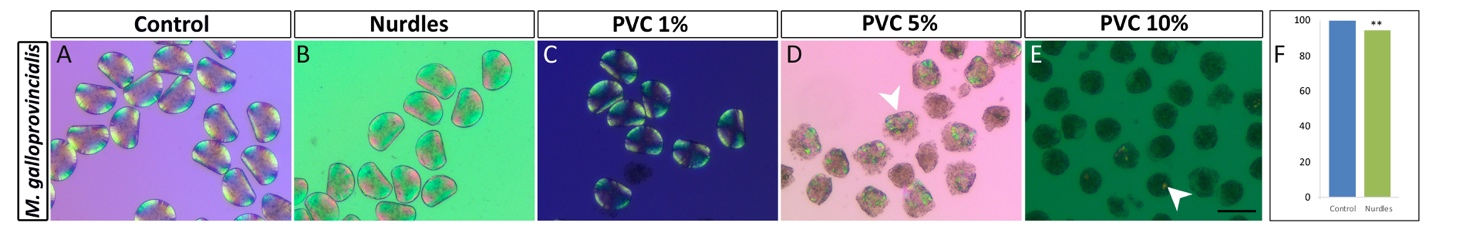

Supplementary Figure 1. Shell formation in *M. galloprovincialis*. Polarised light microscopic images showing the formation of shell in 48 hpf D-larvae. Larve in (A) control, (B) nurdles and (C) 1% PVC exposure conditions show normal shell formation. (D) Larvae exposed to 5% PVC show a smaller shell with a protruding mantle or even an absent shell. (E) In the 10% PVC exposure condition, only a few larvae display some shell specks in the arrested trochophore. Scale bar 100 μm. (F) Comparison between Control D-larvae (blue) and Nurdle leachate D-larvae (green) area size. ** indicate significant differences (unpaired t-test p<0.05). White arrowheads in D and E show examples of incipient shells in each panel.

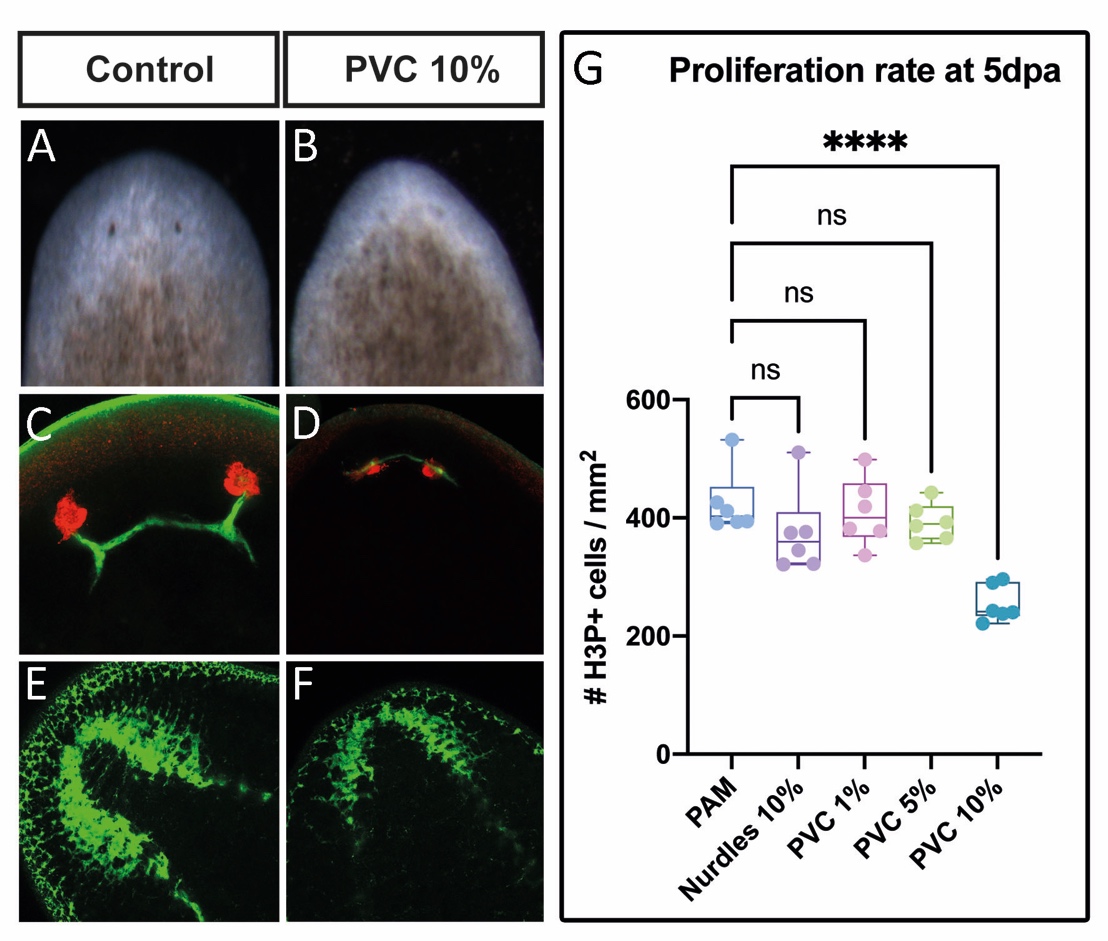

Supplementary Figure 2. Effect of plastic leachates in *S. mediterranea* anterior regeneration. (A-B) Bright field images of 7 days post amputation regenerating heads, and confocal images for (C, D) anti-arrestin (visual system marker) and (E, F) anti-synapsin (panneural marker) immunostainings in 9 days post amputation anterior regenerating heads In C and D the cell bodies of the photoreceptor cells are coloured in red and their axonal projections in green. (G) Proliferation rate in anterior regenerating heads at 5 days post amputation as counts of phosphorylated Histon 3 positive cells per mm^2^. ns indicates no significant differences; **** indicate highly significant differences (p-value 0.0004, Kruskal-Wallis test).
